# Rapid phylogenetic analysis of African swine fever virus from metagenomic sequences

**DOI:** 10.1101/756726

**Authors:** Dongyan Xiong, Xiaoxu Zhang, Junping Yu, Hongping Wei

**Affiliations:** Key Laboratory of Emerging Pathogens and Biosafety, Wuhan Institute of Virology, Chinese Academy of Sciences, Wuhan, China; University of Chinese Academy of Sciences, Beijing, China; African Swine Fever Regional Laboratory of China (Wuhan)

**Keywords:** African swine fever virus, phylogenetic analysis, allele calling, gene-by-gene method

## Abstract

African swine fever virus (ASFV) has devastating impacts on swine health and the world economy. Rapid and accurate phylogenetic analysis of ASFV causing outbreaks is important to reveal diversity and evolutionary of ASFV. Because it is time-consuming and needs biosafety laboratories to isolate ASFV, here we present a new way to perform rapid genome-wide phylogenetic analysis of ASFV using an allele calling based on gene by gene approach directly from genome drafts assembled from metagenomic sequences. Using open-accessed chewBBACA software, 41 publicly available ASFV genomes were analyzed to optimize the parameters and find the alleles. Alleles as many as 94 were found for building the phylogenetic trees, which covered more than 56% of the whole genome. Based on the alleles, current ASFV isolates could be divided into two major clades and a few subclades. Then the method is used to analyze two ASFV genome drafts assembled from two metagenomic sequences of a swine whole blood and a swine spleen tissue collected in Wuhan, China. It shows that the two ASFV genomes showed highest similarity to that of Pig/HLJ/2018 strain and DB/LN/2018 strain, which isolated recently in China. This proved that the ASFV in Wuhan originate from the same source causing the earlier outbreaks in Helongjiang and Liaoning province of China. This method makes it possible to analyze phylogenetic analysis of ASFV draft genomes flexibly without the need of ASFV isolation. Furthermore, because it is based on Allele calling, the ASFV-specific genetic markers found could be translated into clinical diagnostics or can be used broadly to identify conserved putative therapeutic candidates.

## 1. Introduction

African swine fever virus (ASFV) is a contagious virus, which cause lethal ASF disease of wild boars and domestic pigs [1,2]. Since its first outbreak in Kenya in 1921, it has spread to countries in South America, Europe, and Asia [3]. ASFV is a large-encapsulated double-stranded DNA virus with a genome size range from 170 kb to 193 kb, and the genome sizes of different ASFV are quite different. It is currently the only member of the family *Asfarviridae*. It has some similar characteristics to other nuclear DNA large viruses such as *Poxviridae* [4,5]. Neither vaccine nor treatment is available against ASF [6]. In August 2018, African swine fever first broke out in Shenyang, Liaoning Province, China. Then, in a short period of time it spread quickly in China and caused huge economic losses to swine industries [7,8]. Apart from this, Belgium and other European countries had also reported outbreaks of African swine fever since 2018. Interestingly, the reported ASFV in Belgium was quite similar to the previous outbreak in Eastern Europe [9]. Although only 4 ASFV isolates were currently reported, more serious epidemics had occurred in many provinces in China. In August 2019, a new ASF outbroke in small swine farms in Wuhan. Other Asian countries such as Vietnam also discovered the African swine fever epidemic since 2019. As an integrate part of the disease intervention activities, molecular surveillance of ASFV based on phylogenetic analysis methods has played important roles in decision making [10].

At present, there are two main methods for phylogenetic analysis of ASFV. One is based on specific genes or gene segments to construct phylogenetic tree, and the other is genome-based phylogenetic analysis. Most published studies used *B646L* (p72), *E183L* (p54) and *CP204L* (p32) genes or/and central variable region (CVR) of the *B646L* gene for molecular characterization, such as genotyping and subgrouping close-related isolates [11–16]. The phylogenetic analysis using this method is simple, but could only give a rough classification relationship since the coverage of the genes used is about 1% of the whole genome. On the other hand, genome-based approaches are becoming increasingly popular in genomic epidemiology and outbreak detection [17]. For example, David A.G. Chapman et al. performed phylogenetic analysis to show the relationship of Georgia07 isolate with other isolates based on concatenated sequences of 125 conserved ORFs. They also revealed that analysis of *B646L* ORF sequence encoding the p72 major capsid protein does not reflect the phylogeny, because the recombination events may have occurred [3]. Ann Sofie Olesen et al. compared consensus sequence of ASFV Poland15 with Georgia07 focusing on the mutations, indels and SNPs of genome. The result revealed the sequence identity between Poland15 with ASFV Georgia07 [18]. Natalia Mazur-panasiuk et al. performed the global nucleotide genomic alignment to compare seven Polish isolates collected between 2016 and 2017 with other fully annotated genomes falling into various genotypes. The detailed comparative analysis revealed the Polish sequences show the highest similarity to the reference genotype II strain Georgia07 [19]. Xuexia Wen et al. performed a detailed genome comparison of Pig/HLJ/2018 and DB/LN/2018 isolates with the sequences of ASFV-SY18, Georgia07, and other ASFVs reported in European countries in recent years and found that Pig/HLJ/2018 and DB/LN/2018 sequences showed the highest similarity to the sequence of an Poland ASFV isolated in 2017. In addition, ASFV-SY18 is very similar to the sequence of Georgia07 [20]. Jingyue Bao et al. also conducted detailed genomic comparison of China/2018/AnhuiXCGQ and related European p72 genotype II strains. Moreover, the genomes of the ASFV were found containing a Conserved Central Region (CCR) with about 125 kb length, and the phylogenetic analysis were performed based on this region [21]. Zhaozhong Zhu et al. found that there is a large number of indels at both ends of the genome in ASFV leading to a large number of genes inserted or deleted, which contributes to the variety of ASFV genomes. Therefore, the phylogenetic tree of ASFV was constructed based on genomic homologous recombination events [22]. All these methods can describe genomic variations at whole genome level, but is more complicated than the single-gene or gene segments method, requiring isolation of AFSV to get complete genome before analysis. Due to biosafety requirements and slow growth of ASFV, it is usually time-consuming and not convenient to isolate ASFV. In comparison, metagenomic sequencing is much safer and faster to perform by using extracted DNA from samples. Therefore, we intended to develop a method which could be used for phylogenetic analysis of ASFV directly from draft genomes assembled from metagenomic sequences.

In recent years, a method called Core Genome Multilocus Sequence Typing (cgMLST) has been developed for phylogenetic analysis of bacteria [23]. This method is performed by assigning specific alleles to a predefined set of core genes, i.e., genes present in all strains of a given bacterial species. An allele is defined as one of the alternative forms of a mutated gene. A software named chewBBACA is open-available at (https://github.com/INNUENDOCON/chewBBACA), which could perform rapid cgMLST analysis on a laptop computer using Blast Score Ratio (BSR)-Based Allele Calling Algorithm [24]. This software is a complete pipeline for schema creation, validation and allele calling. The first step of the software is using the prodigal software to determine the CDSs in each genome by inputting a set of genomes in FASTA format. Then all the CDSs in the genomes are pairwise-compared in two-steps by removing similar CDSs with smaller length and performing an all-against-all BLASTP search and calculating the BLAST Score Ratio (BSR), resulting a single files containing all unique CDSs in the genomes [25,26]. CDSs with a BSR value greater than 0.6 are considered as alleles of the same locus. After that, the Allele Calling module is used to output a list of CDSs, which could be used in schema evaluation to perform cgMLST/wgMLST by performing multiple sequence alignment using MAFFT [27] and constructing neighbor-joining tree using ClustalW2 [28]. Phylogenetic analysis of bacteria such as *Brucella spp, Yersinia enterocolitica*, etc. has been done successfully [29]. However, no applying this approach has been tried on virus.

In the current study, the chewBBACA approach is used to perform cgMLST of ASFV draft genomes. Phylogenetic analysis of 41 publicly reported ASFV genomes was performed to find the alleles and right parameters first. Then, the method was used successfully for analyzing two ASFV draft genomes. This method provides a flexible and easier way to perform phylogenetic analysis of ASFV without isolation during outbreaks. Furthermore, because it is based on allele calling, the ASFV-specific genetic markers found could be translated into clinical diagnostics or can be used broadly to identify conserved putative therapeutic candidates.

## 2. Materials and Methods

In this study all the analysis were done under a sever with 32G RAM and 2 CPU with 8 core 12 threads on the Linux system.

### 2.1 Phylogenetic analysis using allele calling based gene by gene method

#### 2.1.1 Identification of alleles of ASFV

The software chewBBACA was used to create allelic profile as described previously [24]. Forty-one ASFV complete genomes publicly available were analyzed with ASFV SpainBA71V as the reference genome. The detailed accession numbers of ASFV were available at Supplementary Table S1. The pipline were shown in the flowchart (Figure 1). In order to create suitable allele schema, different -l parameters (100bp, 150bp, 200bp, 250bp, 300bp, 350bp, 400bp, 450bp) were used to remove CDSs with length less than indicated and the -p parameters from 0.8 to 0.99 were set to consider the relatively conserved parts and the more variable features, such as gene deletions among the genomes. Since ASFV uses the same codon usage table with its host, the parameter -ta 1 was chosen during the analysis.

**Figure 1.**
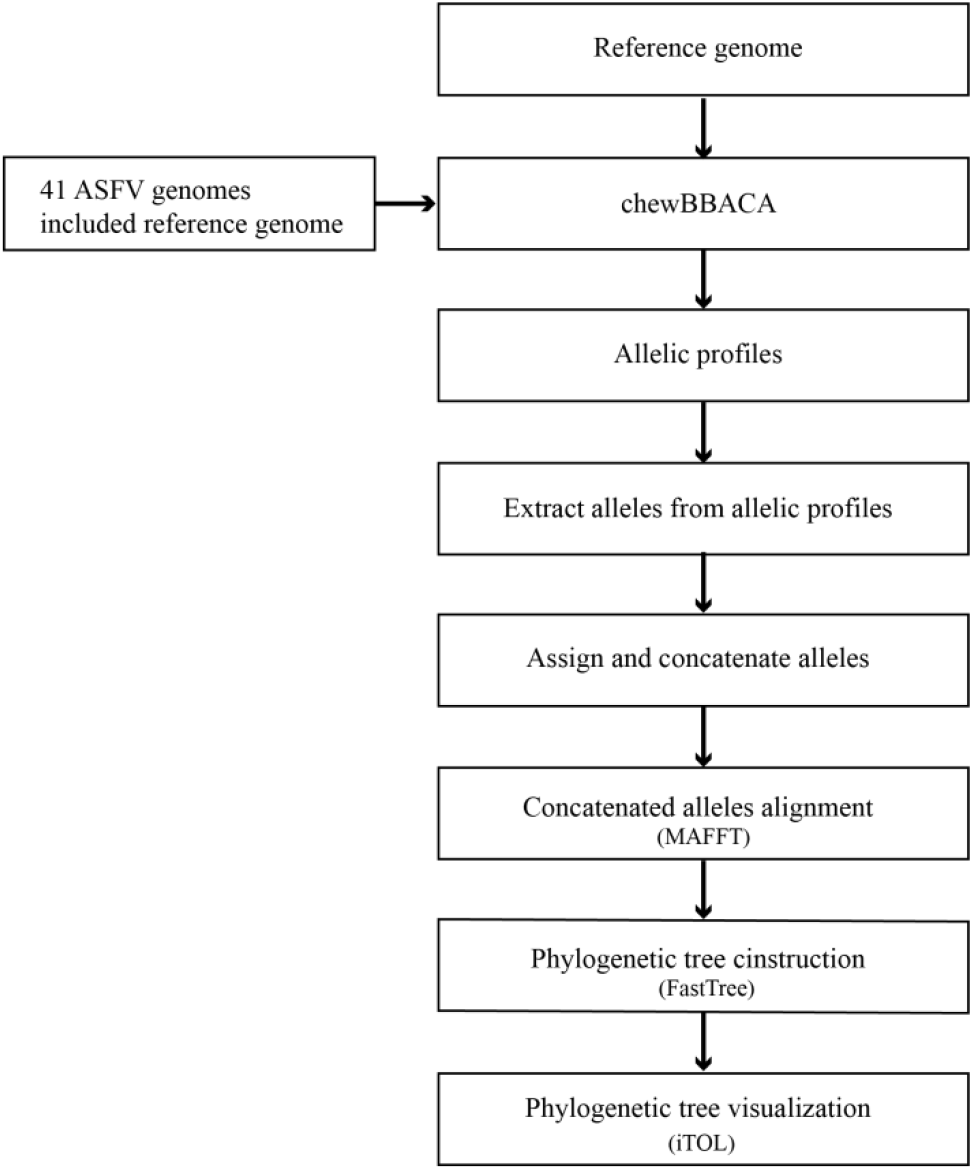
Flowchart for rapid phylogenetic analysis of ASFV

After this step, the chewBBACA outputted allelic profiles containing two important results for the downstream analysis: allelic profile matrix and a set of FASTA files containing the full allelic diversity of each locus.

#### 2.1.2 Assign and concatenate alleles

The alleles of each ASFV genomes were assigned to the respective genomes, then sorted and concatenated under the direction of 5 ‘- 3’ on the genome. All concatenated alleles files were merged into one file.

#### 2.1.3 Sequence alignment and phylogenetic analysis

The nucleotide sequences of the ASFV concatenated alleles were aligned using Mafftv7.4.07 [27] and substitutions, insertions and deletions observed between the aligned sequences were visualized. Phylogenetic relationships among nucleotide sequences were determined by FastTree [32] with the GTR+G and fastest model, each with 1000 bootstrap replicates. The phylogenetic trees were visualized with iTOL [33].

### 2.2 Phylogenetic analysis from metagenomic sequencing data

#### 2.2.1 Sample collection and DNA extraction

In August 2019, outbreaks of African swine fever were suspected in two small swine farms apart about 5 kilometers in Wuhan. Tissues and whole bloods from different pigs were collected. Total DNA was extracted using QIAamp DNA Mini Kit (Qiagen, Hilden, Germany) according to the instruction of the manufacture.

#### 2.2.2 The identification of ASFV through metagenomic sequencing

The total DNA samples were sent to the core facility center, Wuhan institute of virology, CAS for library construction and sequencing. The libraries were constructed using Nextera XT DNA Library Prep Kit (Illumina) and sequenced on Illumina Miseq platform (Illumina, San Diego, USA) utilizing MiSeq^®^ v2 Reagent Kit (Illumina) producing pair-end reads with 250bp length. The FastQC (version 0.11.5) was use to do quality control after the raw reads obtained. The clean reads were obtained by using Trimmomatic-0.36 to filtering the low-quality reads. Then, host genome mapping was done by bwa software (with default options) [30], and all reads mapped to the host genome were removed by samtools (version 1.9, with options -bf 4). The host genome Sus scrofa (version 11.1) is available at ftp://ftp.ensembl.org/pub/release-97/fasta/sus_scrofa/dna/. De novo assembly was executed using SPAdes (version 3.13.0, with the options --meta). Other de novo assembly software such as megahit and SOAP denovo2 were used to correct assembly errors. All the contigs of ASFV were identified by blastn with an ASFV database consisting of the 41 ASFV complete genomes download from NCBI Genbank database [31] and extracted by a Perl script. ASFV contigs assembled by SPAdes under kmer 33 were chosen to conduct phylogenetic analysis using the method described in this study.

### 2.3 Phylogenetic analysis and comparative genomic analysis using complete genomes of Wuhan ASFV

The complete genomes of Wuhan ASFV (GenBank accession numbers: MN393476 and MN393477) were obtained to verify the accuracy of phylogenetic analysis using genome draft. All the ASFV contigs were re-assembled by SeqMan program of LaserGene software and checked based on ASFV reference genomes. Nineteen pairs of primers were designed based on reference genomes to amplify the DNA fragment of two ASFV isolates to fill the gaps and confirm the variable region at both end of the genome. The PCR products were sequenced using the ABI 3730 DNA analyzer (Applied Biosystems™). After the whole complete genome obtained, phylogenetics analysis were also done using the complete genome to check the accuracy of using genome draft. Meanwhile, the pair-wise comparative genomic analysis based on global alignment using the complete genomes was performed by Mafftv7.4.07 to compare the possible variations of Wuhan ASFV strain with ASFV Georgia07, ASFV-Belgium2018/1 as well as ASFV-SY18, China/2018/AnhuiXCGQ and Pig/HLJ/2018 from China, respectively.

In addition, the prediction and annotation of complete genomes of Wuhan ASFV strains were performed using Prodigal and Prokka 1.1.2 software. The putative genes were annotated by BLASTN/BLASTP of annotated genes of ASFV genome.

## 3. Results

### 3.1 Number of alleles found under different parameters

The chewBBACA outputted different number of alleles under different -l and -p parameters. The -1 parameter sets the minimum length of the alleles found, while the -p parameter decides the conservation among the alleles found in all the genomes. As shown in Table 1, a maximum of 94 alleles with -l less than 250 bp and -p 0.80 were found by analyzing the 41 ASFV genomes, while a minimum of 52 alleles were identified with -1 450 bp and -p 0.99.

**Table 1.** Number of alleles identified with different -p and -l parameters

| -l parameter | -p parameter |  |  |  |  |
| --- | --- | --- | --- | --- | --- |
|  | 0.80 | 0.85 | 0.90 | 0.95 | 0.99 |
| 100 bp | 94 | 93 | 91 | 88 | 66 |
| 150 bp | 94 | 93 | 91 | 88 | 66 |
| 200 bp | 94 | 93 | 91 | 88 | 66 |
| 250 bp | 92 | 91 | 89 | 87 | 65 |
| 300 bp | 90 | 89 | 87 | 86 | 64 |
| 350 bp | 87 | 86 | 84 | 84 | 63 |
| 400 bp | 81 | 80 | 78 | 78 | 58 |
| 450 bp | 75 | 74 | 72 | 72 | 52 |

Among the 94 alleles found, 11 genes encode MGFs (MGF 100, MGF 360, MGF 110, MGF 300, MGF 505), which may be related to the pathogenicity of African swine fever [34]; 12 genes encode structural proteins and proteins involved in morphogenesis (*KP177R, A151R, A137R, B438L, B602L, B646L, CP530R, D177L, R298L, E199L, I329L, I267L*) [4,35]; 2 genes encode virulence-related proteins (*A240L, K196R);* 22 genes encode proteins involved in nucleotide metabolism, transcription, replication and repair-related protein (*A859L, F334L* etc.) [4]; 2 genes encode proteins involved host cell interactions (*CP204L, QP383R);* 1 gene encodes anti-apoptotic protein which facilitate production of progeny virions (*A224L*) [36]; 1 gene encodes protein involved in protein regulation (*I215L);* 19 genes encode the putative proteins, such as putative transmembrane protein; and 24 genes encode proteins with unknown functions [21]. The detailed function of these genes are listed in Supplementary Table S2. Besides, these alleles not only existed in CCR region, but also 12 alleles existed outside CCR region, which can provide more extensive information to phylogenetic analysis. Compared with the reported length and position of these genes, the alleles found by chewBBACA are highly consistent.

### 3.2 Phylogentic analysis of ASFV isolates

After adjusting the parameters, we found that it is more reasonable to choose -p parameter from 0.8 to 0.9, -l parameter from 200 to 300, since it can find more conservative alleles with variation information and have more than 56% coverage on the genome after concatenating these alleles. Among the 41 isolates, the genome-wide phylogenetic tree using 91 conserved alleles were mainly grouped into two clades (Figure 2 (a)). Clade 1 includes clade 1.1 containing Kenya 1950 and Kenya 2005 and clade 1.2 consisting of the isolates from Uganda and Kenya 2006. Clade 2 includes clade 2.1 (Malawi 83 only) and clade 2.2, which could be divided into clade 2.2.1 and clade 2.2.2. Clade 2.2.1 includes strains from Malawi, South Africa and Namibia. Clade 2.2.2 is further divided into clade 2.2.2.1 which contains 11 strains in Spain, Portugal, Italy and one strain in South Africa and separates from the rest of the strains, and the clade 2.2.2.2 which includes 13 strains isolated in Europe (Belgium, Poland, Estonia, Georgia, Russia) and 4 strains emerged in China. Interestingly, ASFV strains causing the outbreaks in Belgium and China have a high degree of similarity.

**Figure 2.**
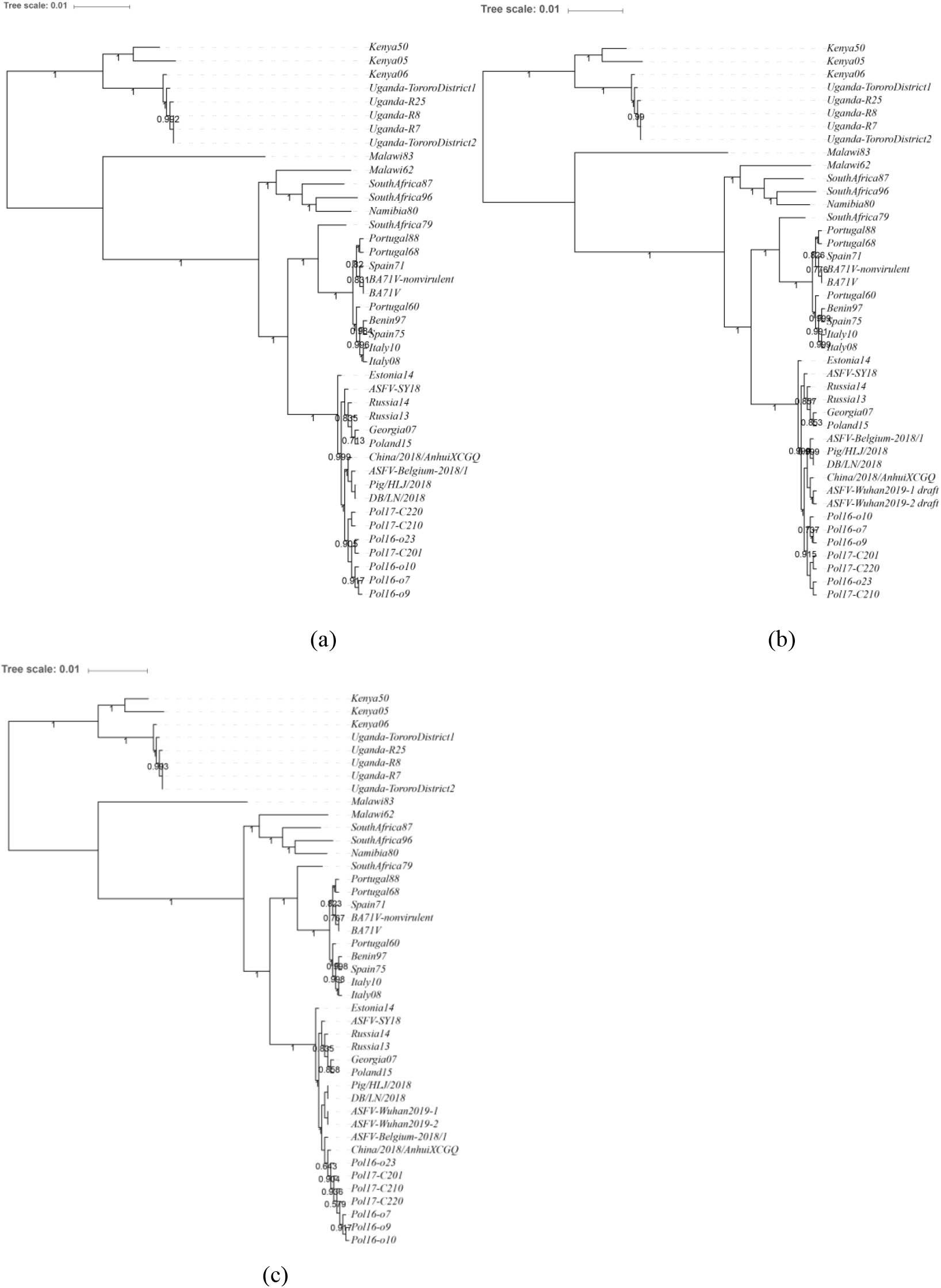
The phylogenetic trees constructed with ASFV isolates by the allele calling based gene by gene method. (a) Concatenate 91 alleles of 41 ASFV public available with the parameter -l 250 and -p 0.85. (b) The allele calling performed using draft genomes of ASFV-Wuhan2019-1 and ASFV-Wuhan2019-2 assmbled by SPAdes software under kmer 33 with the parameter -l 200 and -p 0.9. (c) The phylegenetic tree constructed using complete Wuhan ASFV genomes under the parameter -l 200 and -p 0.9.

### 3.3 Rapid phylogenetic analysis of Wuhan ASFV from metagenomic sequencing

Meanwhile, the draft genomes of two Wuhan ASFV were obtained respectively by SPAdes under the kmer 33 from metagenomic sequencing data. Both of two Wuhan ASFV draft genomes had 35 contigs with length ranging from 200bp to 120kb, which can be obtained at Supplementary File S1. Figure2 (b) showed the classification result using draft genomes of Wuhan ASFV. The Wuhan ASFV isolates were closely related to China/2018/AnhuiXCGQ, Pig/HLJ/2018, DB/LN/2018 and ASFV-Belgium-2018/1.

To verify the accuracy of using genome draft, the two complete genome sequences of Wuhan ASFV isolates were obtained with both 190576bp length and the nucleotide sequences of these two isolates were identical. Figure2 (c) showed the phylogenetic analysis result using whole complete genome. The Wuhan ASFV isolate were found closest to Pig/HLJ/2018 and DB/LN/2018, which agrees with the phylogenetic tree built on the draft genomes.

Further analysis of the ASFV isolates in the clade 2.2.2.2 of phylogenetic tree constructed using complete Wuhan ASFV genomes showed that mutations, insertions as well as deletions happened in each allele provide the basis for classification of the ASFV isolates. As shown in Figure 3, there were many variations in the 36 alleles of ASFV genomes. For example, *A498R, A506R, K145R, NP419L* had more mutations, and *CP204L, QP383R* had more indels. In addition, Estonia14 and Russia14 showed more gene deletions. Within the six Chinese isolates, there was no difference between the alleles of ASFV-Wuhan2019-1 and ASFV-Wuhan2019-2, as well as Pig/HLJ/2018 and DB/LN/2018, but they were different from ASFV-SY18 with 2 mutations and 4 or 5 indels among 36 alleles. Two genes deleted in ASFV-SY18 were *C84L* and *I177L*. Only Pig/HLJ/2018 and DB/LN/2018 have indel deletion in *D1133L* [37]. In addition, among all the strains of clades 2.2.2.2, only ASFV-Belgium-2018/1 had mutations in *D117L*, but it had highest similarity to Chinese isolates. Detailed genetic variation information was shown in Supplementary Table S3.

**Figure 3.**
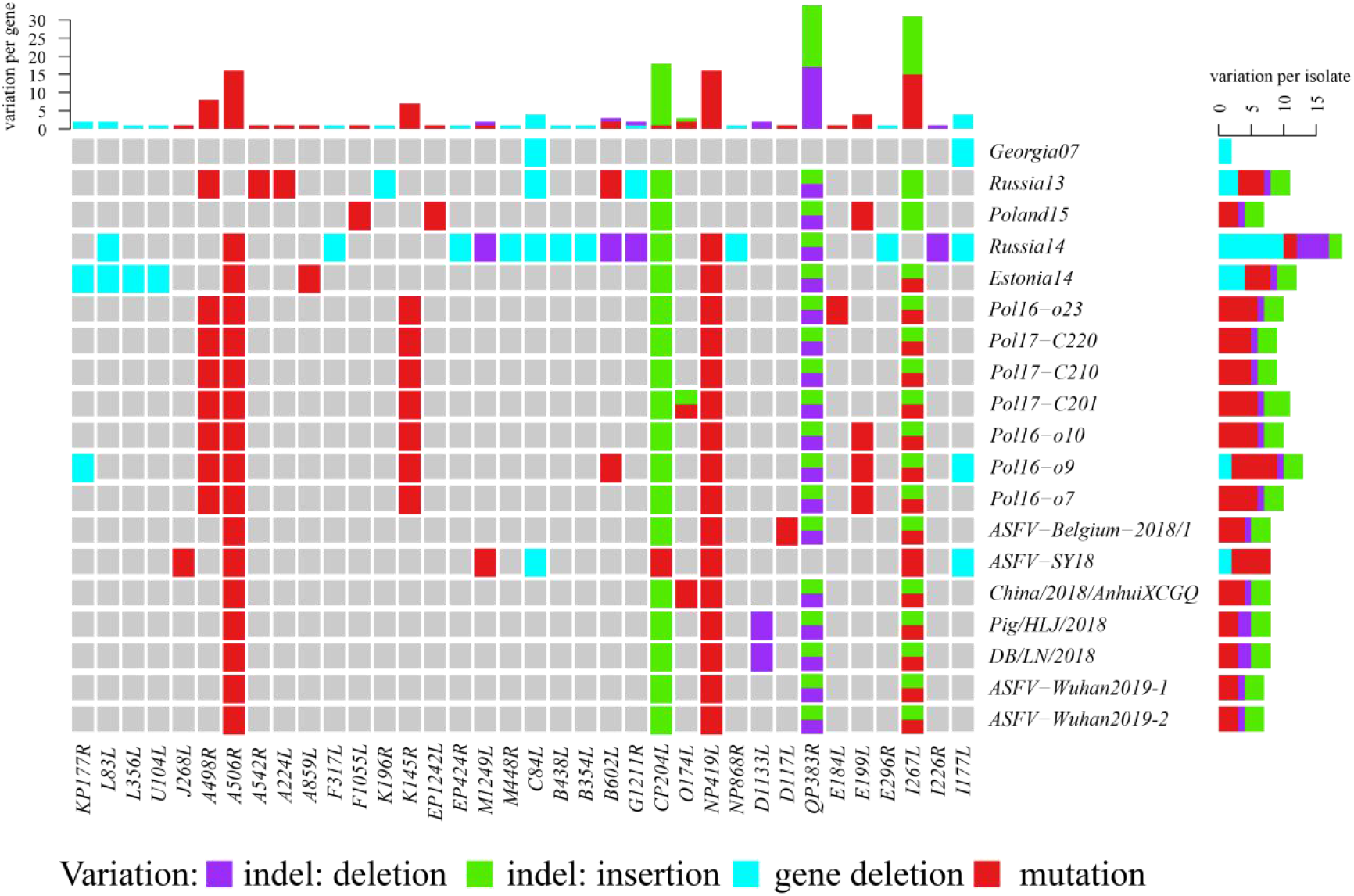
Genetic variations in the conservative genes of ASFV isolates in the Clade2.2.2.2

### 3.4 Pair-wise comparative genomic analysis

From the results of genome-wide analysis, Wuhan ASFV isolates had little variations compared to ASFV Pig/HLJ/2018 and DB/LN/2018. The significance differences between Wuhan ASFV strains and other strains lies in left variable region (LVR) of ASFV genome according to the pair-wise comparative genomic analysis. The Wuhan ASFV strain had a longer genomic segment than the rest of the ASFV strains outbroke in Chinese at LVR of the genome. Besides, ASFV-Belgium-2018/1 strains had a longer genomic segment than all ASFV strains outbroke in Chinese at right variable region (RVR), even if the first outbroke time was so closer between Belgium and China. The other variations contained indels and mutations most occurred in ploy region and non-coding region. The detailed variations of global alignment were shown in Supplementary File S2.

## 4. Discussion

In this work, we present a new way to perform rapid genome-wide phylogenetic analysis of ASFV using an allele calling based on gene by gene approach directly from genome drafts assembled from metagenomic sequences. The gene allele calling algorithm is normally used for bacterial genome analysis.

Although ASFV genome sequences are longer and varied significantly between isolates, 41 published ASFV genomes analyzed by the method showed that alleles as many as 94 could be found and used to construct the phylogenetic trees, which cover more than 56% of the whole genome of ASFV. Therefore, the current method could provide more detailed classification of ASFV isolates than the phylogenetic analysis based single-gene or multiple-genes. As shown in Figure 2, ASFV genomes could be divided into different clades, which were consistent with the present reports by genome-wide analysis [21,38] and the pair-wise genomic alignment analysis in this study.

This approach is also quite flexible by setting different parameters of -l (minimum length of allele) and -p (minimum percentage of loci presence) to identify not only conserved genes but also mutated or deleted genes. Especially, if the -p parameter is chosen to 0.99, the allele genes found are most conservative, which could be considered as target genes for ASFV detection. If the -p parameter is selected below 0.80, alleles with more mutations or indels will be mined, which can help to analyze the genetic variations of different isolates more deeply.

Not only whole genome sequences but also genome drafts could be used to do phylogenetic analysis by this approach [24]. By comparative genomic analysis, the Wuhan ASFV isolates had little variations with two strains Pig/HLJ/2018 and DB/LN/2018 identified in Heilongjiang and Liaoling province and one strain China/2018/AnhuiXCGQ outbroke in Anhui province. Different integrity of genome drafts were assembled from metagenomic data using SPAdes and Megahit under different kmers. The phylogenetic analysis results showed that ASFV strain outbroke in Wuhan clustered together with the strains identified in Heilongjiang, Liaoning and Anhui province as well as the Belgium. Although there is a small difference compared to the result based complete genomes because of incomplete genome assembly under the kmer 33. Thus, the reasonable phylogenetic analysis could be performed using the method we put forward with draft genomes, even if the kmer is as small as 33. Besides, Megahit is the fastest software in most metagenomic denovo assembly programs. Rapid phylogenetic analysis will be done faster from the raw sequencing data using this method consuming less computing resources. Therefore, it could facilitate phylogenetic analysis of ASFV from environmental metagenomic samples, which might be useful for tracing the spread of ASFV in environments.

The recent outbreak of ASF in China had caused significant losses to the industries. Analyzing the genomes of the six isolates in China [20,21] by the current method shows that different from ASFV-SY18, five other isolates (China/2018/AnhuiXCGQ, Pig/HLJ/2018, DB/LN/2018, ASFV-Wuhan-2019-1 and ASFV-Wuhan-2019-2) have a deletion A at the position 728 and a insertion A at the position 762 in *QP383R* gene. These two indels cause twelve amino acids change. *QP383R* encodes NifS-like protein, which has function in host cell interactions. Due to lack of relevant research results, whether the 12 amino acid changes have an effect on the interaction between the virus and the host cells requires further confirmation. Furthermore, China/2018/AnhuiXCGQ has other three mutations from C to T at the position 199, 224 and 328, respectively, in *O174L* gene, causing the changes of the amino acids from P to S, S to F and P to S. Gene *O174L* encodes a repair DNA polymerase, Pol X. This polymerase plays an important role in ASFV base repair with extremely low fidelity and may be involved in strategic mutagenesis of the viral genome [39,40]. These mutations induced three changes happened in the secondary structure of Pol X: Ser 67 being adjacent to αC domain; Phe 75 in the β8 domain; Ser 110 in the αD domain [40]. The mutations may be influence the evolution of ASFV. Besides, only Pig/HLJ/2018 and DB/LN/2018 have deletion A in D1133L gene which encode putative helicases, causing 270nt shortened ORF by frameshift mutation. *D1133L* expressed in the late stage of viral infection [37]. The true function of this gene and whether this change affects the function of gene requires further study. While, ASFV-SY18 has more variations such as two gene deletions of *C84L* and *I177L*, a mutation from C to T at the position 761 causing the change of the amino acid from S to F in the gene *J268L*, a mutation from G to A at the position 364 causing the change of the amino acid from E to K in *M1249L* and a synonymous mutation from G to A at the position 441 and a mutation from A to T at the position 566 causing the change of the amino acid from Y to F in *CP204L* which encodes ASFV-induced protein p32 has function in host cell interactions. In general, some of the variations occurred in functional genes in Chinese ASFV isolates. Moreover, two ASFV discovered in Wuhan have a difference to other Chinese isolates in length at left variation region. Whether these genetic changes of functional genes contribute to epidemiology differences among different provinces in China require further research. It can be speculated that ASFV is changing along with its spread in China, but all of them are close to the isolates in the Europe branch.

Furthermore, we analyzed the species composition from metagenome data, and the result showed the second most abundant virus is porcine type-c oncovirus which is a endogenous retrovirus of pigs. Other bacteria with higher abundance are common bacteria in the farm. But we did not get enough data to do further analysis due to sampling conditions. What can be sure is that the research based metagenome and metatranscriptome sequencing can provide new insight for the prevention and control of ASF epidemic since the cross-infection and interaction between viruses and hosts could be detected.

To be concluded, this study found that the allele calling based gene by gene method could be used for analyzing ASFV phylogenetic relationship from metagenomic sequences rapidly. By setting different parameters of -l and -p, conserved and mutated informations could be obtained from the genomes. In view that chewBBACA could be easily performed on laptops, phylogenetic analysis of ASFV genomes could be done in resources-limited laboratories without ASFV isolation. Besides ASFV, genome-wide analysis of other viruses might be possible using similar approach.

